# The dimerization domain of SARS CoV 2 Nucleocapsid protein is partially disordered as a monomer and forms a high affinity dynamic complex

**DOI:** 10.1101/2024.09.25.614883

**Authors:** Jasmine Cubuk, J. Jeremias Incicco, Kathleen B. Hall, Alex S. Holehouse, Melissa D. Stuchell-Brereton, Andrea Soranno

## Abstract

The SARS-CoV-2 Nucleocapsid (N) is a 419 amino acids protein that drives the compaction and packaging of the viral genome. This compaction is aided not only by protein-RNA interactions, but also by protein-protein interactions that contribute to increasing the valence of the nucleocapsid protein. Here, we focused on quantifying the mechanisms that control dimer formation. Single-molecule Förster Resonance Energy Transfer enabled us to investigate the conformations of the dimerization domain in the context of the full-length protein as well as the energetics associated with dimerization. Under monomeric conditions, we observed significantly expanded configurations of the dimerization domain (compared to the folded dimer structure), which are consistent with a dynamic conformational ensemble. The addition of unlabeled protein stabilizes a folded dimer configuration with a high mean transfer efficiency, in agreement with predictions based on known structures. Dimerization is characterized by a dissociation constant of ∼ 12 nM at 23 ^O^C and is driven by strong enthalpic interactions between the two protein subunits, which originate from the coupled folding and binding. Interestingly, the dimer structure retains some of the conformational heterogeneity of the monomeric units, and the addition of denaturant reveals that the dimer domain can significantly expand before being completely destabilized. Our findings suggest that the inherent flexibility of the monomer form is required to adopt the specific fold of the dimer domain, where the two subunits interlock with one another. We proposed that the retained flexibility of the dimer form may favor the capture and interactions with RNA, and that the temperature dependence of dimerization may explain some of the previous observations regarding the phase separation propensity of the N protein.

---

The SARS-CoV-2 Nucleocapsid (N) protein is responsible for the packaging of the 30 kb-long viral RNA into small viral particles of about 100 nm^1^. The N protein is a flexible and dynamic protein with two structured domains, the RNA binding domain (RBD) and the dimerization domain, flanked by three disordered regions, the N- and C-terminal tails and the linker connecting the RBD and the dimerization domain^2^. Previous investigation of the SARS-CoV N protein pointed to oligomerization as a scaffolding mechanism that favors a dense organization of the RNA genome^3–5^. While oligomerization appears to be key to the function of SARS-CoV-2 N, little is known about the mechanism regulating the assembly of oligomers and how oligomerization impacts the flexible regions of the protein, starting from the dimer assembly. Recent Analytical UltraCentrifugation (AUC) experiments have revealed the formation of stable dimers at concentrations as low as tens of nanomolar^6^. However, AUC measurements at low nanomolar concentrations are challenging, providing only a rough estimate of the dimerization constant. Furthermore, the folded structure of the dimer suggests dimerization most likely occurs as a result of a change in the fold of the protein upon binding. Indeed, the structure of the dimer reveals an N- and a C-terminal 3_10_ helix that encloses a series of short helical segments and two larger beta-strands interspersed by disordered regions^7^ (**Fig. 1**). The binding interface arises from the intertwining of the two molecules, with pairing of the two beta-sheets from each protein subunit and insertion of small helical segments of one subunit into the other^3^ (**Fig. 1**). No structural details are known regarding the conformational changes needed in the monomeric structure for the formation of such a stable dimer. The limited interface and number of contacts within the dimer suggests that the same structure is most likely not stable when the domain is monomeric. Therefore, dimerization must occur through some degree of unfolding (unless the region is already disordered) and refolding of the monomeric unit.

**Figure 1.**
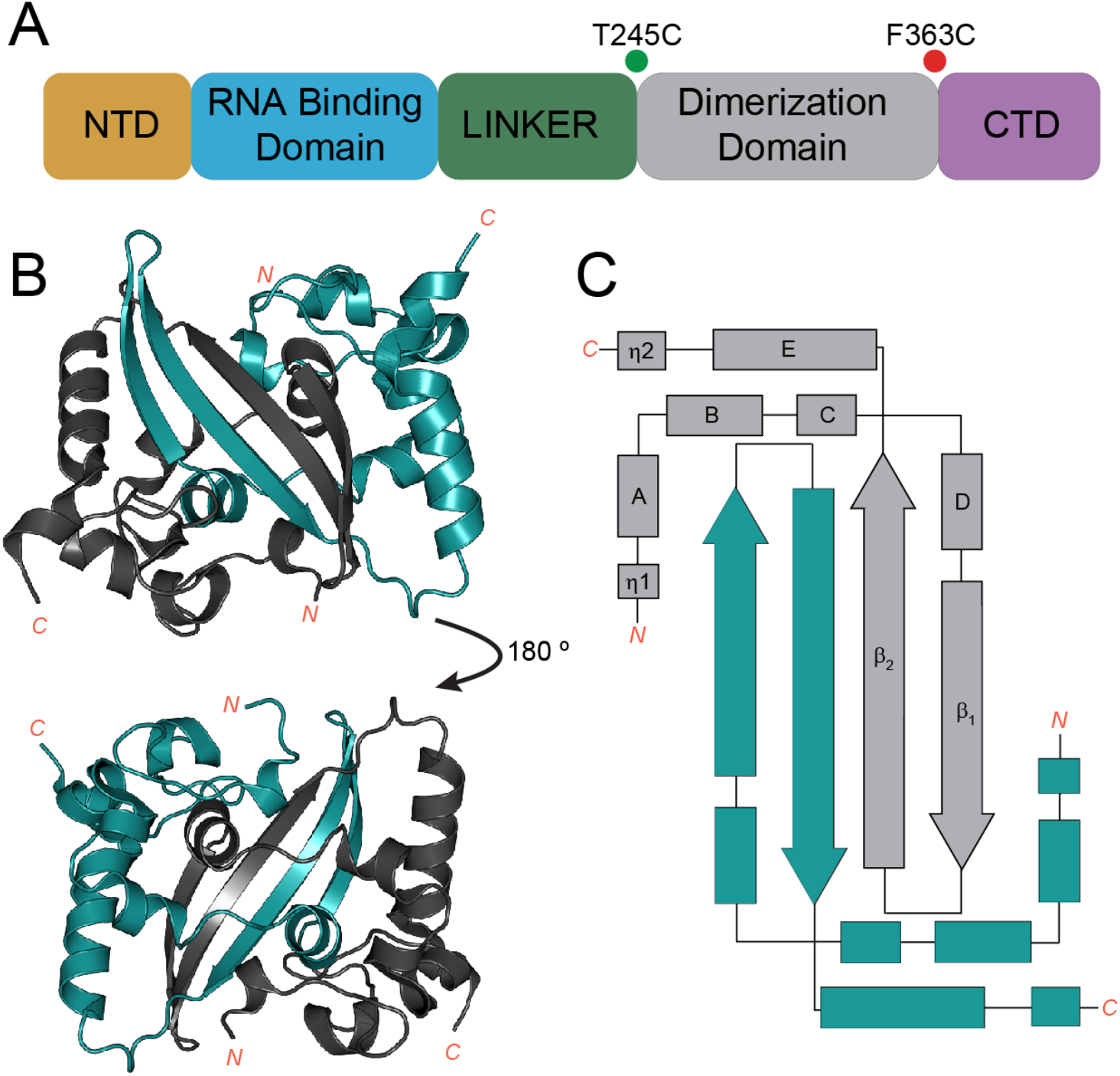
Domain architecture of SARS-CoV-2 N protein and structural features of the dimerization domain. **A.** The SARS CoV 2 N protein contains 5 distinct domains: the N-terminal Domain (NTD), the RNA Binding Domain (RBD), a linker domain (LINK), the Dimerization Domain (DD), and the C-terminal Domain (CTD). **B.** Structure of the Dimerization Domain in dimer form^3^ (PDB: 6WZO) with the two units highlighted in distinct colors: tealand gray. **C.** 2D-topology of the Dimerization Domain when in a stable complex. Letters indicate the main alpha-helices, η_1_ and η_2_ indicate 3_10_ helices, and β_1_ and β_2_ represent beta strands.

Here, we set out to investigate the mechanism of assembly of the dimer and its impact on the other domains of the protein using confocal single-molecule Förster Resonance Energy Transfer (FRET). This approach enables us to monitor the conformations of the full-length protein at sufficiently low concentrations (≤100 pM) to access the monomeric form and to characterize the thermodynamics of dimer formation and the associated conformational changes across the full-length protein. We complemented our observations with all-atom simulations that corroborate the experimental findings and provide atomistic details of the system.

## METHODS

### Single-molecule FRET

The full-length protein variants containing cysteine residues for labeling have been expressed in *E.coli*, purified, and sequentially labeled with Alexa Fluor 488 and 594. Single-molecule measurements have been performed using a modified MT200 instrument (Picoquant, Germany). For all measurements, unless otherwise stated, we used 100 pM of labeled protein (as estimated from serial dilution of a sample of known concentration, based on absorption at 280 nm). Pulsed Interleaved Excitation^8,9^ (PIE) has been used to distinguish the donor-acceptor labeled molecules from donor-donor and acceptor-acceptor species. All measured data were analyzed using the Mathematica package Fretica (https://schuler.bioc.uzh.ch/programs/) to extract mean transfer efficiencies, lifetimes, and labeling stoichiometry ratio, as previously described^2,10,11^.

### All-atom simulations

Simulations for the monomer ensemble were performed as described previously^2^. Briefly, for monomer ensembles, all-atom molecular dynamics simulations were used to generate 200 unique starting conformations for the monomeric dimerization domain^2,12,13^. From those starting structures, 200 independent Monte Carlo simulations were performed in which the partially folded region of the dimerization domain was held fixed while the remainder of the protein was fully sampled^2,14^. This provided an ensemble of 200,000 conformers for further analysis. For dimer ensembles, simulations used the dimeric structure from PDB:6YUN as a starting point, from which the C-terminal IDR was constructed and fully sampled in 39 independent Monte Carlo simulations. This provided a final ensemble of 46,800 conformers for further analysis. All simulation analysis was performed using MDTraj and SOURSOP, and Jupyter notebooks for all analysis, along with the simulation trajectory information, are provided at https://github.com/holehouse-lab/supportingdata/tree/master/2023/cubuk_dimer_202315,16.

Extended Methods are described in the **Supplementary Information**.

## RESULTS

To enable the study of N protein dimerization and associated conformational changes, we designed a full-length N protein construct with cysteine residues at position 245 and 363 (**Fig. 1**) These two residues are expected to be in close proximity upon dimerization based on the known crystal structure^3^. Simulations of transfer efficiency distribution using an AlphaFold^17,18^ prediction of the dimer (the chosen labeling positions extend beyond the residues in the available PDB structures) and FRETraj^19^ (to account for photon statistics) provide an estimate of mean transfer efficiency for this dye pair of approximately 0.90 ± 0.06 (**Fig. S1**). Such close proximity is due to the specific fold of the dimer domain, and we expect the monomer form to adopt more expanded configurations. We will refer to this construct as DD_FL_.

### The Dimerization Domain is partially disordered in the monomer form

As a first step, we investigated the conformations of the DD_FL_ at a concentration of 100 pM. To confirm the protein is monomeric at this concentration, we mixed equimolar concentrations of two single-labeled (F363C) protein preparations that were labeled with either donor or acceptor and observed no appearance of a population with 1:1 donor:acceptor stoichiometry at 100 pM total concentration (50 pM of each species). Under the buffer conditions used in this work (50 mM Tris, pH 7.4, 150 mM KCl), contributions from dimer formation in the stoichiometric plot are observed only above 600 pM of total protein concentration (**Fig. S1**). After confirming that the protein is monomeric at this concentration, we investigated the conformations of DD_FL_ in its monomer form. The histogram of transfer efficiencies reveals a peak centered at 0.567 ± 0.005, which is clearly at odds with the value of ∼0.9 expected from the configuration in the folded structure of the dimer (**Fig. S1**).

To further investigate the existence of stable configurations, we tested the effects of denaturant on the monomeric protein. We observed that the addition of Guanidinium Chloride (GdmCl) leads to a constant shift of the mean transfer efficiency toward lower values, as expected for a protein region that is at least partially (if not completely) unstructured (**Fig. 2**). This conclusion is supported by the investigation of the dependence of the lifetime and transfer efficiency, which provides evidence of dynamics on the nanosecond-microsecond timescale and no significant restrictions to the sampled distance distributions (**Fig. S2**). Therefore, we interpret our results assuming the interdye distance is described by a Gaussian chain distribution with a root-mean-square interdye distance of 5.70 ± 0.02 nm. Using the semi-empirical SAW-*ν* distribution recently proposed by Zheng *et al*.^20^, we obtain a similar root-mean-square distance, 5.4 ± 0.2 nm and a scaling exponent *ν* of 0.48 ± 0.09. We want to emphasize that this does not imply the lack of structure, but that compensatory effects between local structure formation and chain dynamics can give rise to similar statistics of an equivalent completely disordered chain. To gain a better understanding of the molecular configurations of this domain, we turned to all-atom Monte Carlo simulations.

**Figure 2.**
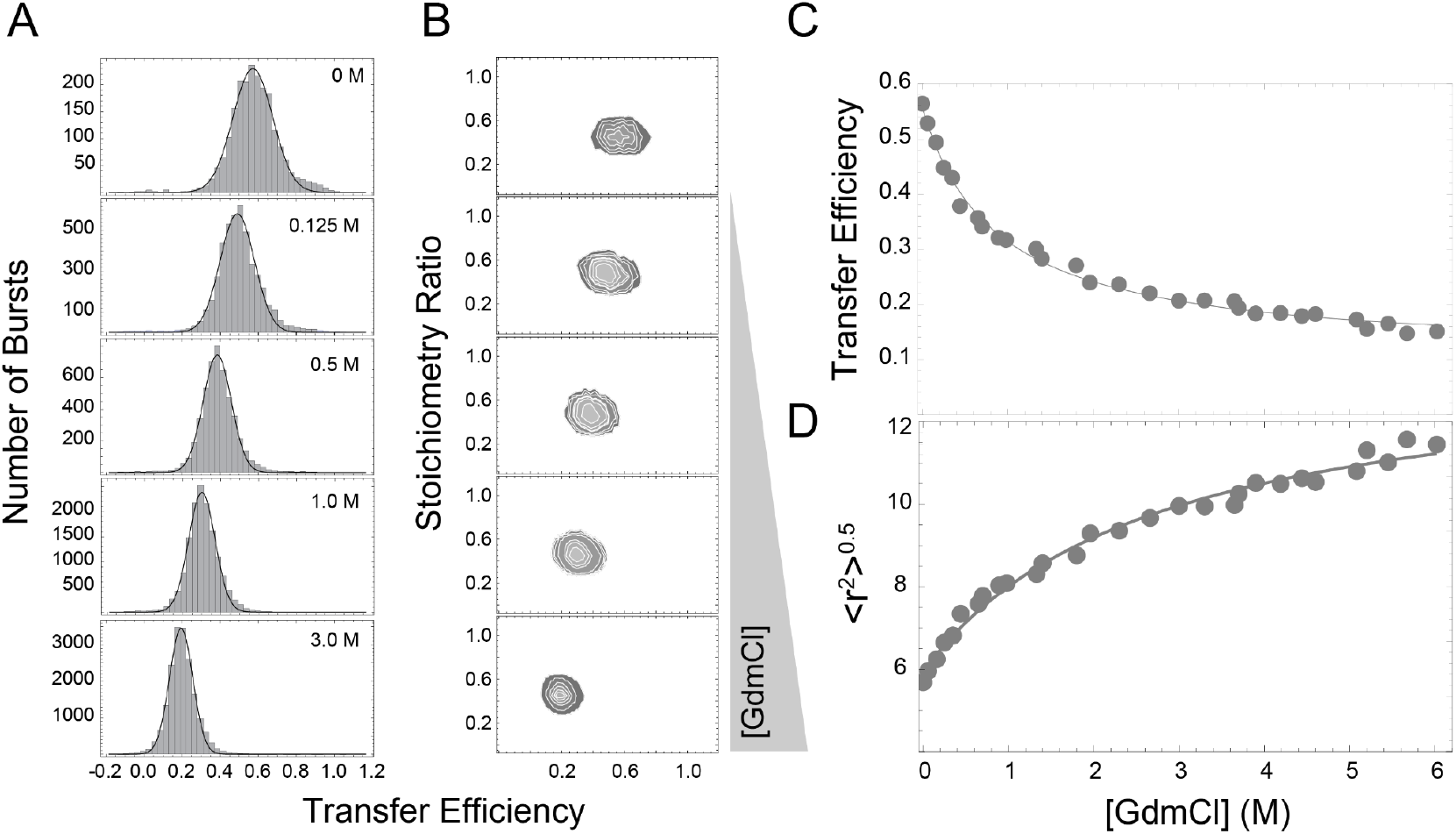
Dimer domain conformations. **A.** Representative histograms of transfer efficiencies for the DD_FL_ construct at 100 pM in aqueous buffer conditions (50 mM Tris, 150 mM KCl) and with increasing denaturant concentrations. **B.** Representative stoichiometry ratio vs. transfer efficiency plots for the corresponding histogram of transfer efficiencies. Stoichiometry ratio of 0.5 indicates 1:1 labeled material, indicating monomeric sample under these conditions. **C.** Distribution of transfer efficiencies as a function of denaturant concentration. **D.** Root-mean-squared interdye distances extracted using a Gaussian chain distribution as a function of denaturant concentration. In panels **C** and **D**, the line represents a fit to the model in **Eq. S7**, which accounts for denaturant binding.

All-atom simulations were performed for a subfragment of the N protein between residues 247 and 419 (**Fig. 3A**). These simulations reveal a root-mean-squared distance between residues 247 and 363 of 4.40 ± 0.04 nm (**Fig. 3B**). While this is smaller than the distance observed from experiments, there are three likely explanations for this discrepancy. Firstly, our simulations lack the extended linker domain, RBD, and NTD, and our previously published single-molecule work and extant SAXS experiments all support a model in which the linker is extended^2,21^ Secondly, our simulations measure the center-of-mass to center-of-mass distance between two residues, a related but distinct measurement compared to the smFRET experiments, which measure the inter-dye distance (where dyes are connected to the protein via short flexible linkers). While a correction factor is used to account for the linker-dye contribution to the distance distribution, deviations may arise because of local excluded volume restriction or interactions of the dyes. Finally, our simulations constrain the configuration of the folded structure in the protein and, as such, may bias the sampled configurations toward more compact ones. With these factors in mind, we take our simulations and experiments to be in reasonable qualitative agreement. In addition to obtaining ensemble average distances, our simulations enable us to calculate complete distributions of intermolecular interactions. The distribution of inter-residue distances between residue 247 and 419 is broad, indicating an underlying ensemble with substantial heterogeneity (**Fig. 3C**). Furthermore, sequence-specific analysis of secondary structure reveals that while partial native secondary structure (native with respect to the dimeric assembly state) is observed for some regions, it is lost in others (**Fig. 3D**). Taken together, our simulations and single-molecule FRET experiments suggest the monomeric form is well-described as a molten globule, with transiently-interacting secondary and tertiary structural elements, but substantial conformational heterogeneity.

**Figure 3.**
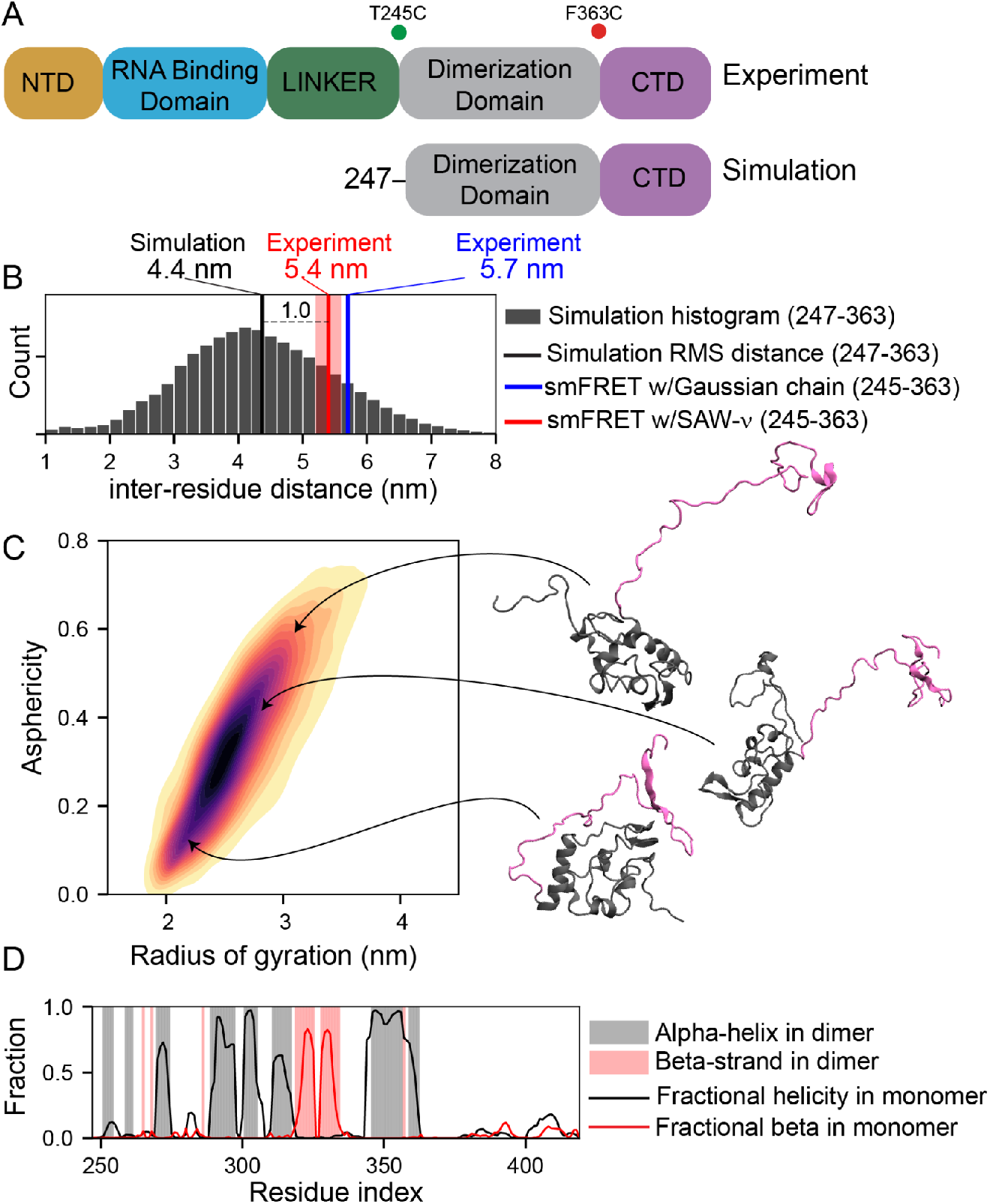
Simulations of dimerization domain. **A.** Schematic showing the domain structure of the full-length construct used in experiments vs. the truncation construct used in simulations. **B.** Inter-residue distance distributions comparing simulations (histogram) and experiment. Lines show the root-mean-squared distance from simulations (black), or the root-mean-squared distances obtained from fitting FRET data to a Gaussian chain model (blue) or the SAW-ν mode (red). **C.** Two-dimensional kernel density estimate showing the distribution of conformational behavior for the monomer ensemble in terms of radius of gyration (ensemble dimensions) and asphericity (ensemble shape). **D.** Overlay of secondary structure propensity comparing monomeric ensemble (lines) vs. dimer-derived structural biases (bars).

### Dimer formation induces folding of the DD domain

We then proceeded to evaluate the protein conformational changes occurring upon dimerization. By titrating increasing concentration of unlabeled protein, we observed the stabilization of a second population at higher transfer efficiency, whose mean transfer efficiency (0.84 ± 0.02) is in good agreement with the expected value based on the folded structure (**Fig. 3A** and **S1**). We then use singular value decomposition (SVD) to quantify the variation of the signal upon binding^22–24^. The advantage of SVD is that it provides a model-free tool to interpret the data without requiring a complex assignment of the distribution of transfer efficiencies to specific conformations of the monomer and dimer^25^. To this end, the measured signal is represented by a matrix **H**, where each line in the matrix refers to a histogram collected at distinct concentrations of unlabeled protein. Singular value decomposition of the matrix **H** is given by

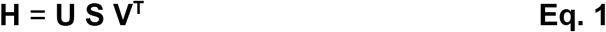

where **S** is the diagonal matrix of the singular values, **U** is the orthonormal matrix of the transfer efficiency distributions of each singular value, and **V^T^** is a matrix containing the amplitude information associated with each concentration of unlabeled protein. SVD can be used to distinguish the contribution of signal and signal changes from experimental noise by comparing the amplitude of the singular values.

As shown in **Fig. 3B**, two major singular values are identified in this titration, whereas all the others contribute to a significantly lesser extent to the total signal. Inspection of the amplitudes further reveals a sigmoidal trend on a logarithmic scale of the concentrations, which reflects the dimerization isotherm of the protein as associated with changes in the first and second singular vector (**Fig. 3D)**. A global fit of the amplitude curves to a binding model that accounts for dimerization of labeled and unlabeled species (**Supplementary Information**) results in a value of the dimerization dissociation constant K_D,L/U_ of 6 ± 2 nM.

### Denaturant effect on dimer stability

We further investigate the stability of the dimer structure in the presence of GdmCl. We chose conditions such that DD_FL_ is complexed in a stable dimer configuration (in the presence of 1 μM unlabeled protein) and then added increasing GdmCl concentrations (up to 1.5 M GdmCl, where the complex is completely destabilized) (**Fig. 4**). We found that with increasing concentration of denaturant (from 0 to 1.3 M GdmCl), the protein adopts more expanded conformations in both the dimer and monomer conformations, with the mean transfer efficiencies of the two states shifting from 0.84 ± 0.02 to 0.70 ± 0.01 (dimer conformation) and 0.567 ± 0.005 to 0.292 ± 0.005 (monomer conformation). Contrasting these observations with the corresponding lifetime information, we confirmed that also the dimer population in the transfer efficiency distribution represents a dynamic conformational ensemble (**Fig. S2**). The corresponding mean interdye distances are reported in **Fig. 4B** and **S4**. In aqueous buffer conditions, we quantified the distance to be 3.8 ± 0.1 nm, according to a Gaussian chain distribution, or 3.7 ± 0.2 nm, according to the SAW-ν model. The scaling exponent for the SAW-ν model is reduced to 0.40 ± 0.02, as implied by the more compact configuration.

**Figure 4.**
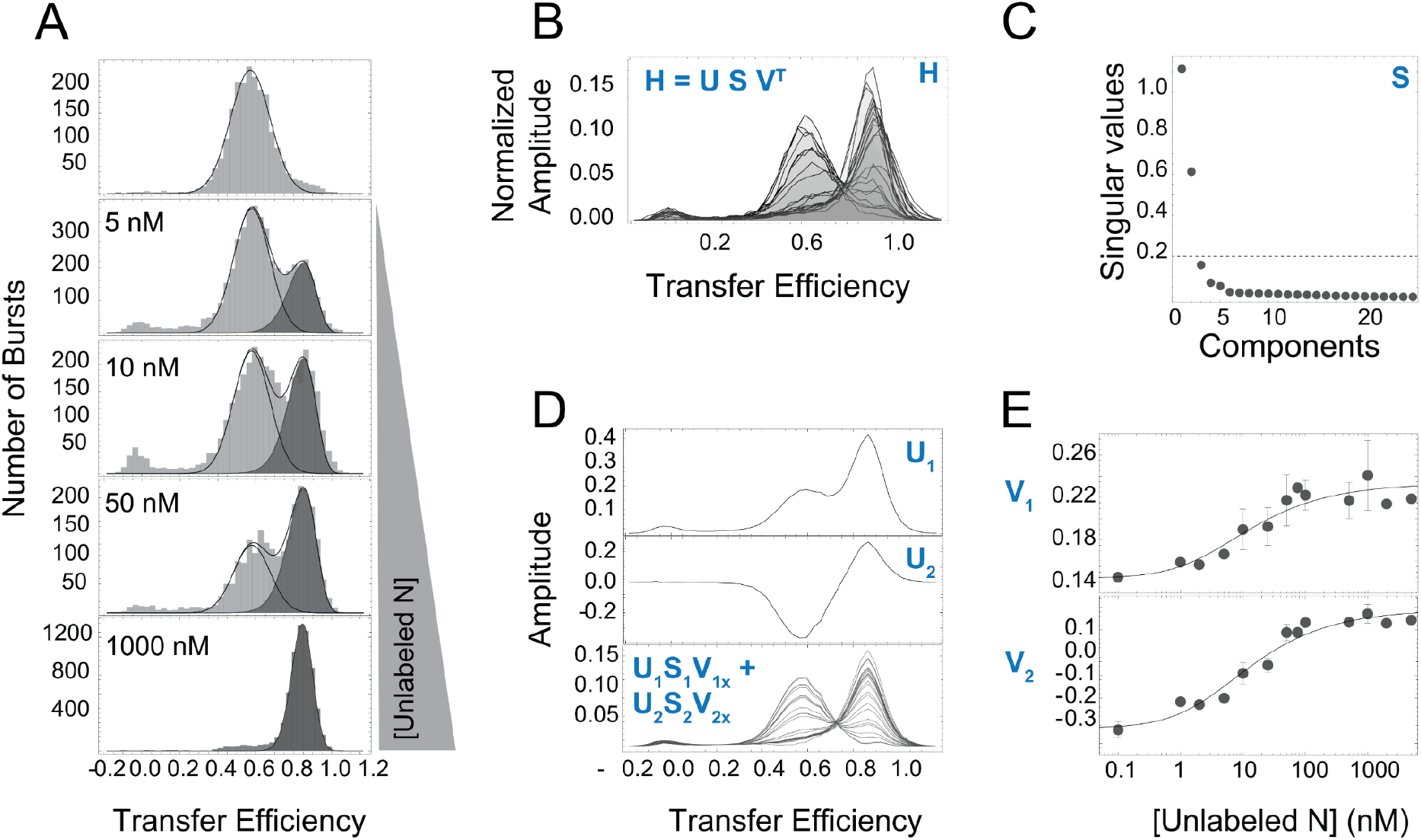
Stabilization of the dimer conformations upon binding: conformational changes and dimerization affinity. **A.** Normalized histograms of transfer efficiencies from 100 pM labeled N protein to a total concentration of 5 μM N protein. **B.** Singular values from the SVD analysis reveal two relevant components. **C.** Basis vectors for each significant singular value. Representative basis vector for component one (top) and representative basis vector for component two (middle). **D.** Reconstruction of histograms using the singular value (S), basis vector (U), and amplitudes (V) (bottom). **E.** Binding isotherms of dimerization of N protein from the SVD analysis of the amplitudes associated with the first (top) and second (bottom) components. Lines represent fit to **Eqs. S9** and **S10**, which account for the binding of the unlabeled N protein to the labeled N protein in solution.

The comparison between lifetime and transfer efficiency is less precise at high transfer efficiencies, making it more difficult to distinguish the effects of chain dynamics from linker dynamics. However, for a completely folded protein, the mean transfer efficiency is expected to remain stable over GdmCl concentration, allowing for changes due to the refractive index^10,26–28^. The expansion of the dimer folded conformation with increasing denaturant concentration (which exceeds the shifts due to refractive index changes and is computed by accounting for such effects) suggests that the observed chain dynamics are not simply the result of dye-linker dynamics. Instead, a certain flexibility must be encoded in the dimer structure, and a certain destabilization of the structure is allowed without losing the stability of the dimer complex. We speculate that these conformational changes may occur along the N- and C-terminal portions of the dimerization domain, which can be destabilized without altering the interface between the intertwining beta-sheets at the core of the dimer structure.

For the monomer, we observe a continuous expansion with increasing concentration of the denaturant, with the protein adopting a root-mean-square radius of 8.5 ± 0.2 nm at 1.3 M GdmCl, which is significantly more expanded than the configuration observed in the absence of denaturant. This conformational change implies that large portions of the structured conformations of the monomeric protein have been destabilized. In addition, the observation of the coexistence of these conformations with a structured dimer implies that, despite some structured conformations having been destabilized and their conformations being expanded, a stable dimer can still be formed. This is not unreasonable, since the formation of the dimer clearly requires large conformational changes in the monomer to allow the two beta sheets to intertwine together.

Analysis of the dissociation constant reveals a linear trend with the denaturant concentration in the regime studied in our single-molecule experiments (**Fig. 4D**). It is interesting to note that the linear trend would suggest a higher fraction of dimer should still be present at 1.5 M GdmCl, but we do not observe such a population. We speculate that this represents a threshold concentration over which either the dimer domain is completely destabilized, or any residual structure in the monomer required for dimer formation has been destabilized (based on equilibrium data we cannot distinguish the two case scenarios). To confirm this hypothesis, we further tested the fraction of dimer at 1.75 M GdmCl with a concentration of protein 10 μM and at 2 M GdmCl at a concentration of 45 μM. According to the linear fit, we should observe a fraction bound of 0.5 and 0.5, respectively, as determined from the linear extrapolation of the 𝑙𝑛 𝐾_𝐷_ versus GdmCl concentration. We found that no dimer complex is formed at any of these conditions, supporting our hypothesis that 1.5 M GdmCl is suppressing dimer formation by denaturing essential folded structures in either the monomer or the dimer.

### Temperature effect on dimer stability

We further investigated the temperature dependence of the dimer stability to quantify the enthalpic and entropic contributions at play and the corresponding conformational changes. To this end, we used a temperature-controlled cuvette, analogous to previous studies^29–31^. To limit any contributions from the pH dependence on temperature, we performed all temperature measurements in the HEPES buffer. Similar to Tris buffer, the protein adopts a transfer efficiency of 0.57 ± 0.03 in the monomer configuration and of 0.85 ± 0.03 in the dimer configuration.

We started by studying the temperature response of DD_FL_ under monomeric conditions to establish a baseline for the conformational changes of this region. We found that the domain adopts more compact conformations as we increase temperature, in line with previous temperature dependence experiments on disordered proteins ^29,30,32^. We then studied the effects of temperature on the domain when the protein is a stable dimer (1 μM). We observed that from 10 °C to 30 °C, the protein remains dimeric and the mean transfer efficiency associated with the conformations of the dimer species report a small shift toward higher values, indicating a small, but measurable compaction of the protein. Starting from 37 °C, a broadening of the distribution of transfer efficiencies is observed. This broader distribution of transfer efficiencies can be disentangled into two populations that represent the monomer and dimer species. Mean transfer efficiencies and corresponding areas of each subpopulation can be used to estimate the association constant at each temperature (**Fig. 5**). When increasing temperature to 50 °C or higher, the protein is completely dissociated and adopts the conformations of the monomeric form.

**Figure 5:**
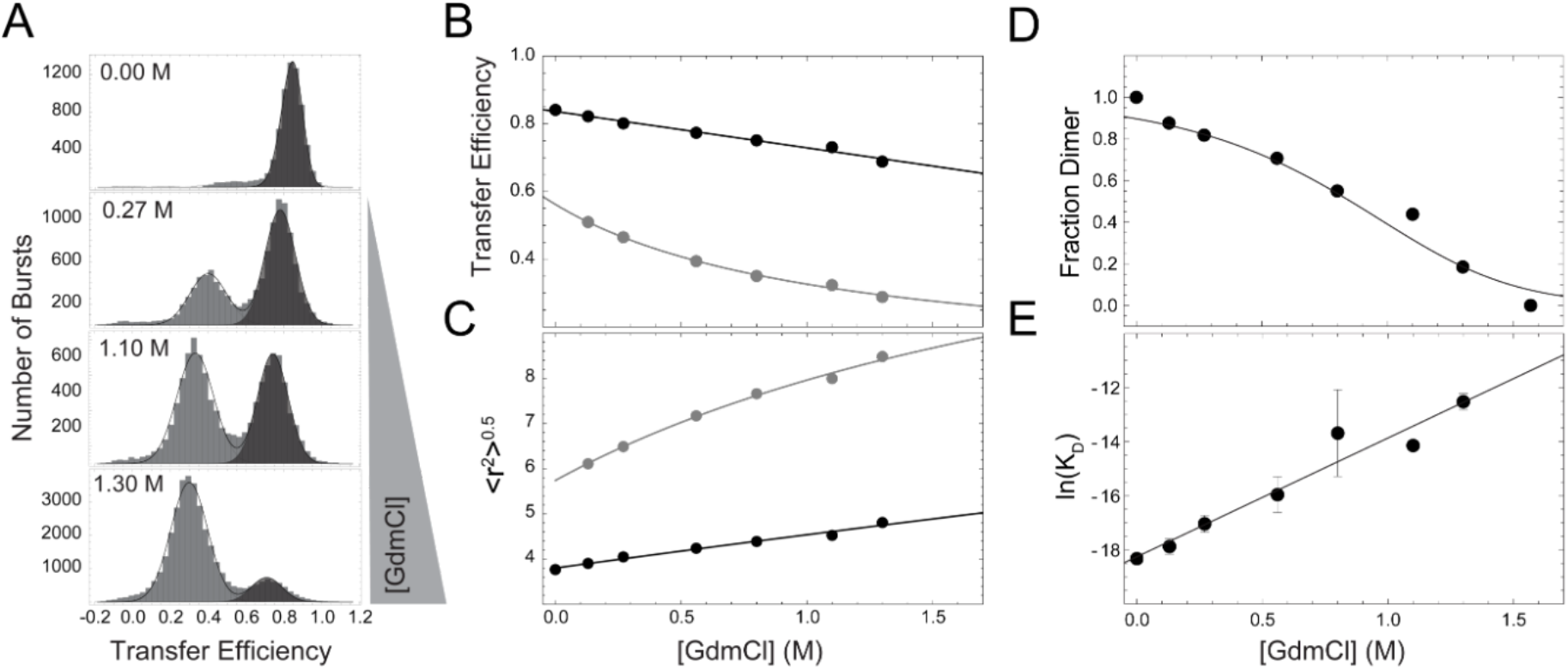
Denaturant dependence of the dimerization domain. **A.** Representative histogram of transfer efficiencies for the DD_FL_ construct in the presence of 1 μM unlabeled protein in the absence and with increasing concentrations of denaturant. **B.** Transfer Efficiencies as a function of denaturant for the dimer (darker gray) and monomer (lighter gray) distributions. Lines represent a fit to the Schellman weak binding model in **Eq. S7**. **C.** Root-mean-squared inter-dye distances as a function of denaturant for the dimer (darker gray) and monomer (lighter gray) distributions. Lines represent a fit to **Eq. S7**. **D.** Plot of the fraction of dimer as a function of denaturant concentration. Here, dimers can be seen present up to 1.3M GdmCl. **E.** Plot of ln(K_D_), with K_D_ expressed in M units, as a function of denaturant. A linear trend is observed between the dissociation constant and the concentration of GdmCl.

**Figure 6:**
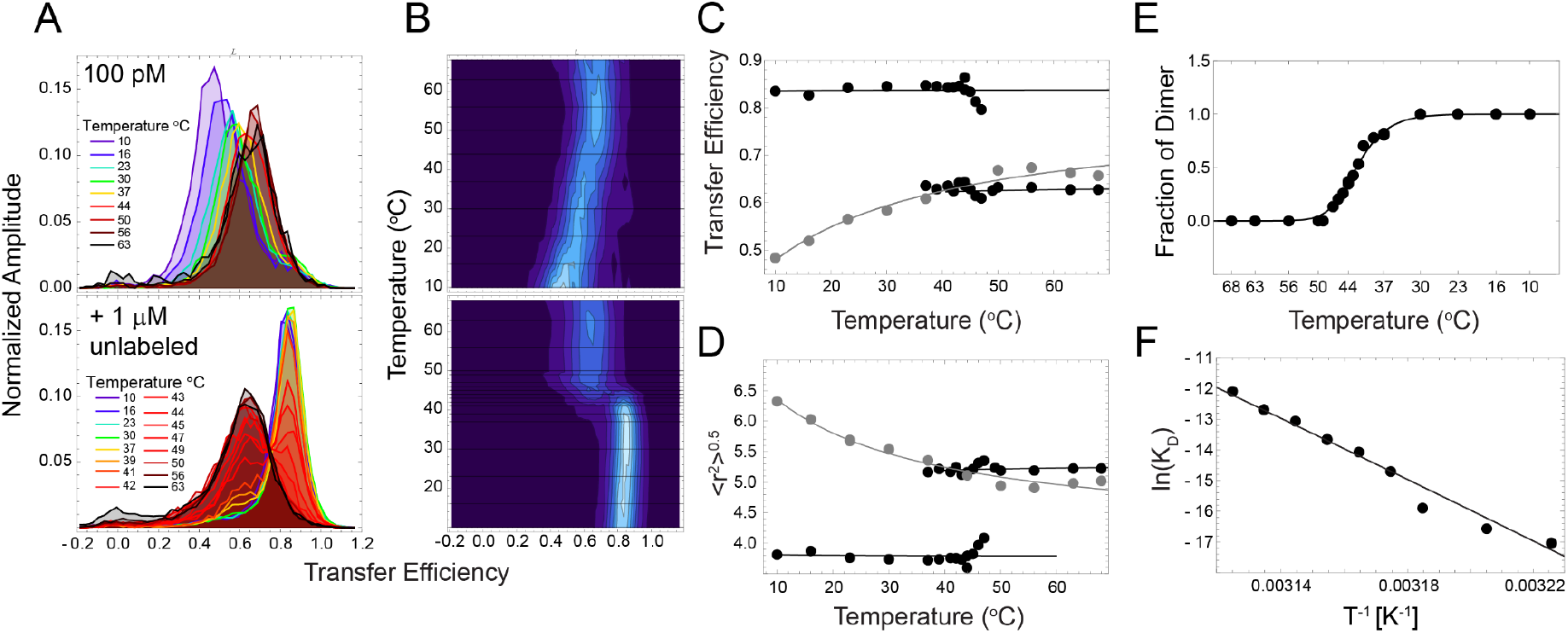
Temperature dependence of the dimerization domain. **A.** Normalized histograms of transfer efficiencies for the DD_FL_ construct at 100 pM (top) and in the presence of 1 μM unlabeled protein (bottom), ranging from 10 °C to 63 °C. **B.** Corresponding 2D Temperature vs. transfer efficiency plots for 100 pM (top) and with the addition of 1 μM unlabeled protein (bottom). **C.** Dependence of transfer efficiency on temperature for 100 pM total protein concentration (gray) compared to 1 μM total protein concentration (black). **D.** Dependence of root-mean-squared interdye distance on temperature for 100pM total protein concentration (gray) compared with 1 μM total protein concentration (black). **E.** Fraction of labeled protein in dimers (formed with unlabeled protein) as a function of temperature. The line is computed assuming the linear dependence of ln(K_D_) as a function of 1/T (K^-1^) shown in panel F, according to **Eqs. S9-11**. **F.** Plot of the ln(K_D_), with K_D_ expressed in M units, as a function of 1/T (K^-1^). The linear fit reports on the enthalpy (slope) and entropy (intercept) of dimer dissociation according to **Eq. 2a**. Values obtained from the fitting procedures are reported in **Supplementary Table 5**.

To analyze the enthalpic and entropic contributions of dimerization, we performed a Van’t Hoff analysis on the temperature dependence of the dissociation constant (**Fig. 5E**). According to Van’t Hoff equation, if in a given range of temperatures the enthalpy of a reaction does not change, the logarithm of the dissociation constant 𝐾_𝐷_ is a linear function of the reciprocal of absolute temperature *T*, where the slope reports about the enthalpy Δ𝐻^0^ and the intercept reports on the entropy Δ𝑆^0^ at standard conditions (1 M) of binding:

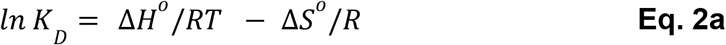

where *R* is the gas constant.

We observed a linear decrease of 𝑙𝑛 𝐾_𝐷_ as a function of 1/*T*, in the range of temperatures from 37 °C to 47 °C. We found that the dimer formation is exothermic, with a ΔH of -99 ± 6 kcal/mol and a ΔS of 0.29 ± 0.02 kcal/mol K. The large enthalpy is compatible with previous estimates for another coronavirus dimerization domain^33^ and likely reflects the coupled folding and binding of the domain.

When extending the plot to lower temperatures and including the 𝐾_𝐷_ determined at 23 °C, we clearly observe a deviation from linearity that suggests a non-negligible contribution of the heat capacity:

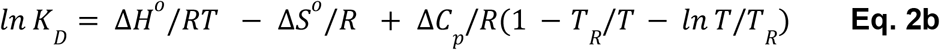

Where 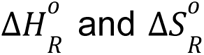 are the enthalpy and entropy at the reference temperature, Δ𝐶𝑝 is the heat capacity, 𝑇 is temperature, and 𝑇_𝑅_ is the reference temperature. Corresponding values of the fit are reported in **Supplementary** Fig. 4 and **Table 5**. Analysis of dimer fractions as estimated from the extrapolation at lower temperatures with **Eq. 2a** and **Eq. 2b** reveals a small discrepancy with respect to the experimentally determined fractions at lower temperatures. We attribute this discrepancy to the contribution of the effect of temperature on the conformations of the dimer domain, which may result in a variation of enthalpic and entropic contributions.

### Does dimerization affect other domains?

We tested whether the formation of the dimer complex induces conformational changes in the other N protein domains (**Fig. S5**). To this end, we compared the conformations of the N-terminal domain (NTD), Linker, and C-terminal domain (CTD), with and without saturating concentrations of unlabeled proteins. At 150 mM KCl, the conformations of each region of the protein match our previous observations, where all three populations exhibit a narrow distribution of transfer efficiencies^2^. Upon addition of 1 μM of unlabeled protein, we observed no significant shift in the NTD (with mean transfer efficiency shifting from 0.60 ± 0.01 to 0.595 ± 0.003) or RBD (with mean transfer efficiency shifting from 0.818 ± 0.003 to 0.828 ± 0.004). We detected a small expansion of the LINK (with mean transfer efficiency shifting from 0.570 ± 0.003 to 0.520 ± 0.004) and a significant expansion of the CTD (mean transfer efficiency shifting from 0.64 ± 0.02 to 0.570 ± 0.001). The conformational changes of the Linker and of CTD are consistent with the suppression of previously identified interactions between the DD domain and surrounding disordered regions^2^.

### Conformational ensemble of the dimerization domain and C-terminal IDR

Finally, we examined intra- and inter-molecular interactions of the C-terminal IDR in the dimer context using all-atom simulations. We performed simulations in which the dimerization domains were held in place to match the previously published crystal structure, but the C-terminal tails (residues 362 – 419) of each protomer were fully sampled (**Fig. S6A**). These simulations reveal a root-mean-square distance between residues 247 and 363 of 3.0 nm (**Fig. S6B and C**). As with the monomeric construct, the inter-residue distance is somewhat more compact than the values obtained by experiment, but given the absence of dimerization domain flexibility or an account of the dye linkers, these results are in reasonable agreement. The protomers in the dimer are more compact and less heterogeneous when compared to the monomeric dimerization domain in isolation (**Fig. S6B**). Moreover, these simulations suggest the C-terminal IDRs interact minimally with one another but do interact with surface residues on the folded dimerization domain (both in *cis* and in *trans*), predominantly via a cluster of hydrophobic and negatively-charged residues between positions 393 and 403 (**Fig. SD, E**). In short, our simulations of the dimeric construct are consistent with experiments and suggest folding leads to a reduction in dimerization domain dimensions and heterogeneity, yet the C-terminal IDR remains highly disordered, even in the dimeric state.

## Discussion

Our experiments and simulations provide new insights into the conformational properties of the dimerization domain in the context of the full-length protein, both in its monomer and dimer forms, and on the energetics associated with dimerization.

An approximate estimate of the dissociation constant was previously determined using Analytical Ultracentrifugation^6^. In the work of Zhao *et al.*^6^, upon dilution, the appearance of a monomer population was observed between 30 nM and 1 nM, resulting in an approximate K_D_ of roughly 30 nM for unlabeled protein and 2 nM for labeled protein. Our single molecule approach enabled us to follow the transition from monomer to dimer with a fine-tuned titration, starting from 100 pM (labeled concentration) and increasing concentration up to 10 μM (using unlabeled protein). We modeled the data accounting for the formation of dimers between labeled and unlabeled proteins (labeled-labeled, unlabeled-unlabeled, and labeled-unlabeled) and the corresponding impact on the association constants (**Supplementary Information**)^34^. Following this procedure, our experiments yield a dissociation constant between labeled and unlabeled molecules K_D,L/U_ of 6 ± 2 nM and a dissociation constant between unlabeled (or labeled) molecules K_D,U/U_ of 11 ± 3 nM. Our results are in reasonable agreement with the previous estimates reported by the Schuck group^35^. The good agreement between these results suggests we are capturing the same phenomenon, besides small differences that can be introduced by solution conditions and specific labeling positions of the protein.

In addition to quantifying the dimer dissociation constant, our experiments enabled access to conformational changes in the monomer and dimer forms of N protein. We found that in the monomer form, the dimerization domain of the N protein exhibits a significant degree of flexibility and is very sensitive to solution conditions (as probed by denaturant and temperature). We reason that this flexibility may be required for facilitating the formation of the specific fold of the dimer domain, where beta sheets belonging to the two proteins are required to intertwine with each other. The dimer form, while much more compact, also exhibits a dynamic behavior and can be modulated by solution conditions. In particular, denaturants can significantly expand the conformations of the complex.

We speculate that the high stability of the complex and its inherent flexibility can be harnessed by the nucleocapsid protein to enhance the ability to recruit and trap the viral RNA. Indeed, a flexible chain ensures a larger captured radius than a conventional folded domain^36^. At the same time, dimerization ensures a higher valence of interaction, which can increase the effective affinity for the nucleic acid. We note that our results are likely applicable to a series of analogous dimerization domains in other coronaviruses^7,37^ (SARS-CoV^4^, hCoV-NL63^38^, MERS-CoV^39^, MHV-A59^40^, IBV^41^), since these all share a common fold. While similar with respect to the structure adopted by the dimer domain, the sequence composition differs across viruses and may modulate thermal stability, affinity, as well as conformational heterogeneity of monomeric and dimeric forms. Understanding how the sequence modulates the energetics and conformational properties of the dimer may provide insights into future emerging coronavirus and help design drugs that target dimer stability (limiting the valence of the N protein) or its conformational flexibility (limiting the capture radius and adaptable interface for binding RNA).

Our experiments also provide a possible molecular mechanism of the lower critical solution temperature (LCST) previously identified in phase separation experiments with the N protein. Previously it has been shown that, at room temperature, N protein requires RNA to undergo phase separation *in vitro* ^2,42^. While the exact critical temperature was not identified, further work has established that N protein also undergoes phase separation without RNA starting from a temperature of 45 °C at protein concentrations between 1 and 4 μM^43^. This range of temperature coincides with our temperature-dependent suppression of protein dimerization at the same protein concentration and is consistent with independent differential scanning fluorimetry measurements^37^. We reason that the unfolding of the dimer domain and the large flexibility and disorder of the monomer can augment the multivalence of N protein and favor its phase separation and propensity to aggregate in the absence of RNA. Interestingly, in support of this hypothesis, we detect small variations in the mean transfer efficiency of the monomer population of the dimerization domain in presence of micromolar concentrations of unlabeled N protein. This discrepancy observed above 37 °C differs from what observed in presence of denaturant at 23 °C under similar protein concentrations. These observations suggest that at sufficiently high temperatures the monomeric form of the protein starts to form homotypic interactions that do not lead to the known-fold of the dimer.

Interaction with RNA complicates this scenario, introducing a new set of interactions between the protein and the nucleic acid (e.g., through the RNA binding domain), which can be realized through both disordered and structured domains. While it was suggested that the binding of RNA could destabilize the dimer structure^43^, we do not observe any alteration of the binding fraction when titrating non-specific single-stranded RNA as well as specific double-stranded RNA (**Fig. S4**). In light of our experiments, the expansion of the dimerization domain upon binding of RNA, as observed in cryo-EM measurements^37^, may reflect the inherent flexibility of this dimer complex and its adaptability for favoring RNA binding. Future work is required to carefully evaluate how the multivalence of RNA interactions rewire the protein-RNA phase diagram, its temperature dependence, and how these interactions control the condensation of single and multiple RNA chains.

## Conclusions

Here, we have completed the characterization of the conformational properties of the N protein by investigating the structural ensemble of the dimerization domain. We have found that the dimerization domain forms a high-affinity complex (starting from low nanomolar concentrations) that retains part of the flexibility intrinsic to the monomeric form. Our results pave the way to constructing quantitative models of the protein-protein and protein-RNA interactions at play when the N protein condenses viral genomic RNA or undergoes phase separation.

## Contributions

J.C. performed and analyzed experiments. ASH performed and analyzed simulations. JJI provided research tools. KH, MDSB, and AS supervised the work. JC, MDSB, and AS designed the experiments. JC, JJI, KH, ASH, MDSB, and AS have written the manuscript.

## Acknowledgements

We thank Upasana L. Mallimadugula for her valuable input on figures and Debjit Roy for help with the temperature setup. This research was supported by the NIH National Institute on Allergic and Infectious Diseases with R01AI163142 (to A.S., K.B.H., and A.S.H.). The content is solely the responsibility of the authors and does not necessarily represent the official views of the NIH.

