## Supplementary material for "The dimerization domain of SARS CoV 2 Nucleocapsid protein is partially disordered as a monomer and forms a high affinity dynamic complex": SI

### Supplementary Information.

**Protein expression, purification, and labeling.** GST-$His_9$-SARS-CoV2 WT (cysless), NTD-FL, RBD-FL, LINK-FL, and CTD-FL were expressed and purified as previously described[2].

GST-$His_9$-SARS-CoV2 $DD_{FL}$ Nucleocapsid protein was expressed recombinantly in Gold BL21(DE3) cells (Agilent). 6 L cultures were grown in LB medium with carbenicillin (100 µg/mL) to $OD_{600}$ ~ 0.8 and induced with 0.25 mM IPTG for 3 hours at 37ºC. Harvested cells were lysed with sonication at 4ºC in lysis buffer (50 mM Tris pH 8, 300 mM NaCl, 10% glycerol, 10 mg/mL lysozyme, 5 mM βME, cOmplete™ EDTA-free Protease Inhibitor Cocktail (Roche), DNAse I (NEB), RNAse H (NEB)). The supernatant was cleared by centrifugation (140,000 x g for 1.5 hr) and the pellet was resuspended in 50 mM Tris pH 8, 300 mM NaCl, 10% glycerol, 6 M Urea, 5 mM βME and incubated at 4ºC for one hour. The resuspension was cleared by centrifugation (140,000 x g for 1 hr), and the GST-$His_9$-N protein in the supernatant was bound to a FF HisTrap column (GE Healthcare) in buffer A (50 mM Tris pH 8, 300 mM NaCl, 10% glycerol, 20 mM imidazole, 5 mM βME) containing 6 M Urea. The column was then washed with buffer A allowing the protein to refold on the column. The column was then washed with high salt buffer (Buffer A +2 M NaCl), followed by Buffer A. The GST-$His_9$-N protein fusion was then eluted with buffer B (buffer A containing 500 mM imidazole) and dialyzed into cleavage buffer (50 mM Tris pH8, 50 mM NaCl, 10% glycerol, 1 mM DTT) containing HRV 3C protease. FL N protein was then bound to an SP sepharose FF column (GE Healthcare) and eluted using a gradient of 0-100% buffer B (buffer A: 50 mM Tris pH 8, 50 mM NaCl, 10% glycerol, 5 mM βME, buffer B: buffer A + 1 M NaCl) over 120 min. Purified N protein was analyzed using SDS-PAGE. Protein concentrations of stock solutions were determined spectroscopically in 50 mM Tris (pH 8.0), 200-500 mM NaCl, 10% (v/v) glycerol using extinction coefficients of 42530 $M^{-1}$ $cm^{-1}$(FL).

**Protein labeling.** All Nucleocapsid variants were labeled with Alexa Fluor 488 maleimide (Molecular Probes) under denaturing conditions in buffer A (50 mM Tris pH 8, 6M Urea) at a dye/protein molar ratio of 0.7/1 for 2 hrs at room temperature. Single labeled protein was isolated via ion-exchange chromatography (Mono S 5/50 GL, GE Healthcare - protein bound in buffer A(+5 mM BME) and eluted with 0-40% buffer B (buffer A + 1 M NaCl) gradient over 70 min) and UV-Vis spectroscopic analysis to identify fractions with 1:1 dye:protein labeling. Single labeled Alexa Fluor 488 maleimide labeled N protein was then subsequently labeled with Alexa Fluor 594 maleimide at a dye/protein molar ratio of 1.3/1 for 2 hrs at room temperature. Double labeled (488:594) protein was then further purified via ion-exchange chromatography (Mono S 5/50 GL, GE Healthcare).

**Experimental setup and procedure for single-molecule fluorescence experiments.** Single-molecule experiments were conducted on a Picoquant MT200 instrument (Picoquant, Germany). Pulsed Interleaved Excitation (PIE) was realized by alternating the pulses from a diode laser (LDH-D-C-485, PicoQuant, Germany) and a supercontinuum laser (SuperK Extreme, NKT Photonics, Denmark) filtered by a z582/15 band pass filter (Chroma). The excitation rate was set to 20 MHz such that a delay of approximately 25 ns occurs between each of the two laser pulses.

Laser beams were focused through a 60x1.2 UPlanSApo Superapochromat water immersion objective (Olympus, Japan), and emitted photons were collected through the same objective. Emitted photons were further filtered through a dichroic mirror (ZT568rpc, Chroma, USA), a long pass filter (HQ500LP, Chroma Technology), and the confocal pinhole (100 μm diameter) before being selected according to polarization and emission wavelength. Donor and acceptor photons were first separated by a dichroic mirror (585DCXR, Chroma) and further refined by using band pass filters, ET525/50m or HQ642/80m (Chroma Technology), respectively. Finally, the filtered photons were focused on SPAD detectors (Excelitas, USA), and their arrival time was recorded with a HydraHarp 400 TCSPC module (PicoQuant, Germany). All FRET experiments used a donor excitation with a laser power of 100 μW (measured at the back aperture of the objective) and an acceptor excitation adjusted to match the total emission intensity after donor excitation (between 50 and 70 μW).

All measurements, unless differently specified, were performed in 50 mM Tris pH 7.4, 200 mM β-mercaptoethanol (for photoprotection), 0.001% Tween20 (for surface passivation), and GdmCl at the reported concentrations. Measurements were performed in uncoated polymer coverslip cuvettes (Ibidi, Wisconsin, USA) and pegylated coverslips (for temperature dependence). When using denaturant or salt, the exact concentration is determined by measurements of the solution refractive index with an Abbe refractometer (Bausch & Lomb, USA). Each sample was measured for at least 10 min at room temperature (295 ± 0.5 K).

**Construction of transfer efficiency histograms.** Photons were time-binned in bins of 1 ms. Contiguous bins with at least one bin above 15 photons were merged into a single burst, and bursts with at least 20 photons were selected for further analysis. The exact threshold was selected based on the background contribution identified in the photon counting histograms with 1 ms binning. Transfer efficiencies for each burst were calculated according to

$$E = n_A/(n_A + n_D) \quad \textbf{Eq. S1}$$

where $n_A$ and $n_D$ are the numbers of donor and acceptor photons, respectively.

Transfer efficiencies were corrected for background, acceptor direct excitation, channel crosstalk, differences in detector efficiencies, and quantum yields of the dyes.

The labeling stoichiometry ratio S was computed accordingly to:

$$S = I_D/(I_D + \gamma_{PIE} I_A) \quad \textbf{Eq. S2}$$

where $I_D$ and $I_A$ represent the total intensities observed after donor and acceptor excitation, $\gamma_{PIE}$ is a correction factor accounting for differences in donor and acceptor detection efficiency and laser intensities. In the histograms, we present the bursts with stoichiometry corresponding to 1:1 donor:acceptor labeling (in contrast to donor and acceptor-only populations), which are selected according to the criterion 0.3 < $S$ < 0.7. Variations in the selection criteria for the stoichiometry ratio do not significantly impact the observed mean transfer efficiency (within experimental errors).

**Fit of transfer efficiency distributions.** To estimate the mean transfer efficiency and extract multiple populations from the transfer efficiency histograms, each population was approximated with either a Gaussian or a LogNormal distribution function. When fitting more than one peak, the histogram is analyzed with a sum of Gaussian and/or LogNormal functions.

**Root-mean-square interdye distance.** Conversion of mean transfer efficiencies to an interdye distances were performed after verifying the fast rearrangement of the ensemble based on the comparison of lifetime vs. transfer efficiency. Under this assumption, the conversion is obtained by integrating the distance dependence of the transfer efficiency

$$E(R) = \frac{R_0^6}{R_0^6 + R^6} \quad \textbf{Eq. S3}$$

on the distance distribution according to:

$$\overline{E} = \int_0^\infty E(R)\, P(R)\, dR \quad \textbf{Eq. S4}$$

where $R_0$ is the Förster radius.

For a Gaussian chain, the distribution is given by:

$$P(R) = 4\pi R^2 \left(\frac{3}{2\pi r^2}\right)^{3/2} exp\left(\frac{-3R^2}{2r^2}\right) \quad \textbf{Eq. S5}$$

where $r$ represents the root-mean-square interdye distance of the protein and is the only fitting parameter.

For the SAW-v model can be expressed as:

$$P_{SAW}(R, v) = A_1 \frac{4\pi}{b_0 N^v} \left(\frac{R}{b_0 N^v}\right)^{2+g} Exp\left[- A_2 \left(\frac{R}{b_0 N^v}\right)^{\delta}\right] \quad \textbf{Eq. S6}$$

where $A_1 = \frac{\delta}{4\pi} \frac{\Gamma[5+g/\delta]^{\frac{3+g}{2}}}{\Gamma[3+g/\delta]^{\frac{5+g}{2}}}$, $A_2 = \left(\frac{\Gamma[5+g/\delta]}{\Gamma[3+g/\delta]}\right)^{\frac{\delta}{2}}$, $g = \frac{(\gamma-1)}{\nu}$,$\delta = \frac{1}{(1-\nu)}$, $\gamma = 1.1615$, $\Gamma$is the Euler Gamma Function, $b_0$ = 0.55 nm is an empirical prefactor [20] , *N* is the number of residues between the fluorophores, and $\boldsymbol{\nu}$ is the scaling exponent. Here $\boldsymbol{\nu}$ is the only fitting parameter.

**Binding of Denaturant.** Similar to previous works[2,44], we model expansion of the disordered chain with denaturant with an empirical model based on the denaturant dependence of the Schellman weak-binding model[45]:

$$r(c) = r_0(1 + \rho \frac{K\,c}{1+Kc}) \quad \textbf{Eq. S7}$$

where $r_0$ is the root-mean-square interdye distance in the absence of denaturant, $\rho$ is a term that represents the chain expansion in the presence of denaturant compared to the absence ($r_0$), $K$ is the binding constant and $c$ is the concentration of denaturants. This semi-empirical model is used only for comparison between different titrations, without aiming to extract significant quantities. A more precise description would require the use of a coil-to-globule model, which provides access to the free energy change in the protein chain and can be directly related to the Schellman model[46].

**Fluorescence lifetime.** Fluorescence lifetimes were estimated from bursts using a maximum likelihood method[4748] and implemented in the Fretica package. The dynamic regime was computed according to:

$$\tau_D/\tau_{D0} = 1 - \overline{E} + \frac{\sigma^2}{1-\overline{E}} \quad \textbf{Eq. S8a}$$

$$(\tau_A - \tau_{A0})/\tau_{D0} = 1 - \bar{E} + \frac{\sigma^2}{\bar{E}}$$ **Eq. S8b**

where $\tau_D$ and $\tau_{D0}$ are the donor lifetimes in the presence and absence of the acceptor, $\tau_D$ and $\tau_{D0}$ are the acceptor lifetimes in presence and absence of the donor, $\bar{E}$ is the mean value of the transfer efficiency according to **Eq. S4** and σ represents the corresponding variance, $\sigma^2 = \int E^2(r)P(r)\,dr - \bar{E}^2$ [49]. In the absence of fluctuations over a distribution of distance (i.e., $\sigma^2 = 0$), the equations reduces to the well-known static regime equation where:

$$\tau_D/\tau_{D0} = (\tau_A - \tau_{A0})/\tau_{D0} = 1 - E$$ **Eq. S8c**

**Modeling dimerization.** Binding experiments have been analyzed accounting for the possibility of forming dimers of labeled molecules $DD_L$ with other labeled molecules ($DD_L$:$DD_L$), labeled molecules with other unlabeled molecules $DD_U$ ($DD_L$:$DD_U$), as well as between unlabeled molecules ($DD_U$:$DD_U$). To this end, we imposed a mass balance such that:

$$[DD_L]^{tot} = [DD_L] + 2[DD_L:DD_L] + [DD_L:DD_U]$$ **Eq. S9a**

$$[DD_U]^{tot} = [DD_U] + 2[DD_U:DD_U] + [DD_L:DD_U]$$ **Eq. S9b**

For each dimer we can write the corresponding dissociation constant as:

$$K_D^{LL} = \frac{[DD_L]^2}{[DD_L:DD_L]}$$ **Eq. S10a**

$$K_D^{UU} = \frac{[DD_U]^2}{[DD_U{:}DD_U]}$$ **Eq. S10b**

$$K_D^{LU} = \frac{[DD_L]\,[DD_U]}{[DD_L{:}DD_U]}$$ **Eq. S10c**

Finally, we accounted for the fact that if expressed in the microscopic rate constants of binding and unbinding the $K_D^{LU}$=1/2 $K_D^{UU}$=1/2 $K_D^{LL}$, under the assumptions that the rates of unlabeled and labeled are equal[34].

The fraction of labeled protein that forms dimers with unlabeled protein (coinciding with the FRET population at 0.5 stoichiometry ratio) is given by:

$$f_{Dimer}^{LU} = \frac{[DD_U]^{tot}\left(\left(K_D^{LU}\right)+2\left([DD_L]^{tot}+[DD_U]^{tot}\right)-\sqrt{K_D^{LU}}\sqrt{K_D^{LU}+4\left([DD_L]^{tot}+[DD_U]^{tot}\right)}\right)}{2\left([DD_L]^{tot}+[DD_U]^{tot}\right)^2}$$ **Eq. S11**

**All-atom simulations.**

Simulations for the monomer ensemble were performed as described previously[2]. Briefly, initial all-atom molecular dynamics simulations were performed on the dimerization domain (residues 247–360) using Gromacs 2019 with the AMBER03 force field and explicit TIP3P solvent[13,50–52]. The exploration of conformational space was enhanced using the FAST algorithm[12]. Following these simulations, 200 conformationally-distinct structures were identified as starting states for the monomeric dimerization domain. Using these 200 starting structures, subsequent all-atom Monte Carlo simulations were performed in which the backbone dihedral angles of the dimerization domain residues (268 – 360) were frozen while other regions (247 – 268, and 361 – 419) were fully sampled. All sidechain dihedrals for all residues were fully sampled. Monte Carlo simulations were performed using the ABSINTH implicit solvent model and CAMPARI simulation engine (https://campari.sourceforge.net/) [14]. Each individual simulation was run at 330 K for 24 x $10^6$ Monte Carlo steps, with the first 4 x $10^6$ steps discarded for equilibration. Post equilibration, conformations were saved every 20 x $10^3$ Monte Carlo steps. In all, our simulations generated an ensemble of 200,000 conformers from 200 independent simulations. Simulations were analyzed using SOURSOP and MDTraj [15,16].

For simulations of the dimeric protein, the dimer structure from PDB:6YUN was used as a starting structure[37]. The structure encompasses residues 249 to 364. Using this dimeric structure as a starting state, residues 249–361 in both protomers had their backbone dihedral angles held fixed, and the remaining residues (362 – 419) were constructed as random non-overlapping starting configurations in both protomers. Finally, the promoters were held together by distance restraints, ensuring the dimer remained fixed in its starting orientation. Following structure preparation, 39 independent all-atom Monte Carlo simulations were run in which the C-terminal tail (362 – 419) was fully sampled, whereas only sidechain dihedrals in the folded domains were sampled. Each simulation was performed for 64 x $10^6$ Monte Carlo steps, with the first 4 x $10^6$ steps discarded as equilibration and conformers saved every 50 x $10^6$ Monte Carlo steps. In total, simulations yielded a dimeric ensemble of 46,800 conformers from 39 independent simulations.

#### Supplementary Figures

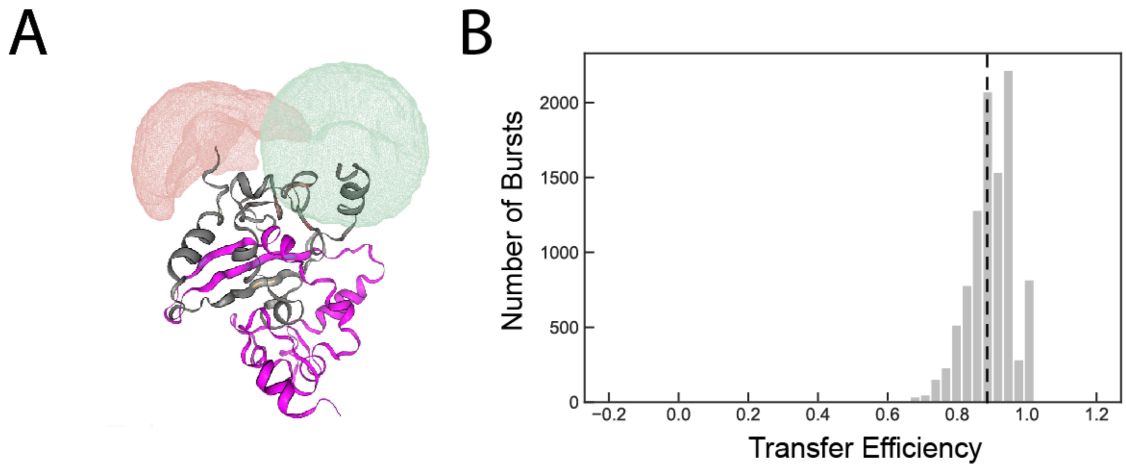


**Supplementary Figure 1. Estimated transfer efficiency based on the dimer structure. A.** Structure estimated using AlphaFold with labeling positions 245 and and 363. Dye-clouds based on available excluded volume for the dyes computed using FRETraj. **B.** Corresponding distribution of transfer efficiencies generated using FRETTraj.

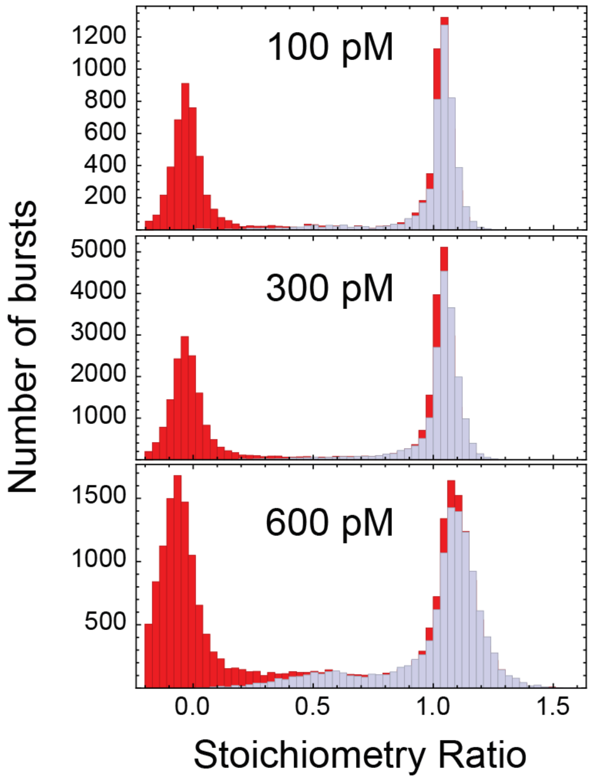


**Supplementary Figure 2.** Labeling Stoichiometry Ratio plots of $DD_{FL}$ measured at 100 pM, 300 pM, and 600 pM total concentrations of nucleocapsid protein. Single-labeled F363C Alexa-488 was mixed with Single-labeled F363C Alexa-594 at equimolar concentrations. We do not observe any population at 0.5 stoichiometry under 600 pM, which would represent the formation of a dimer.

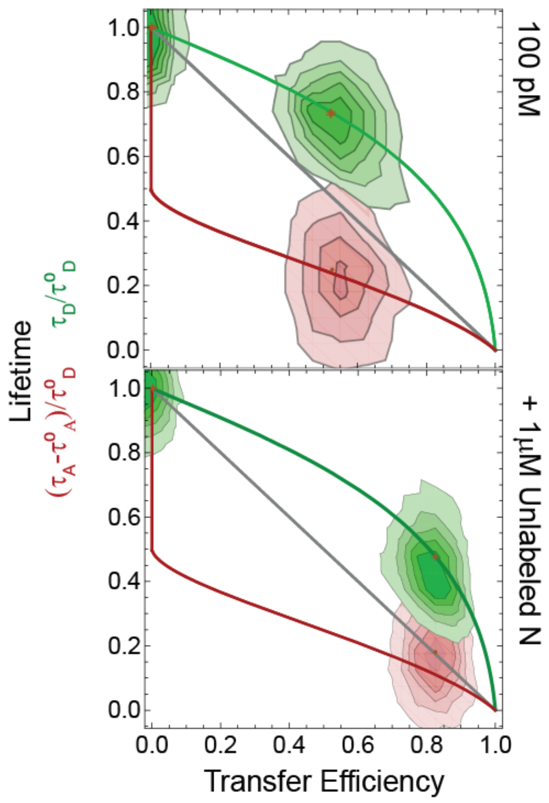


**Supplementary Figure 3.** Lifetime vs Transfer efficiency of the $DD_{FL}$ at 100pM (top) and with the addition of 1 μM unlabeled N protein (bottom). In both cases, the populations sit near the dynamic line (green for donor, red for acceptor), as opposed to falling on the static line (gray), indicating a rigid configuration.

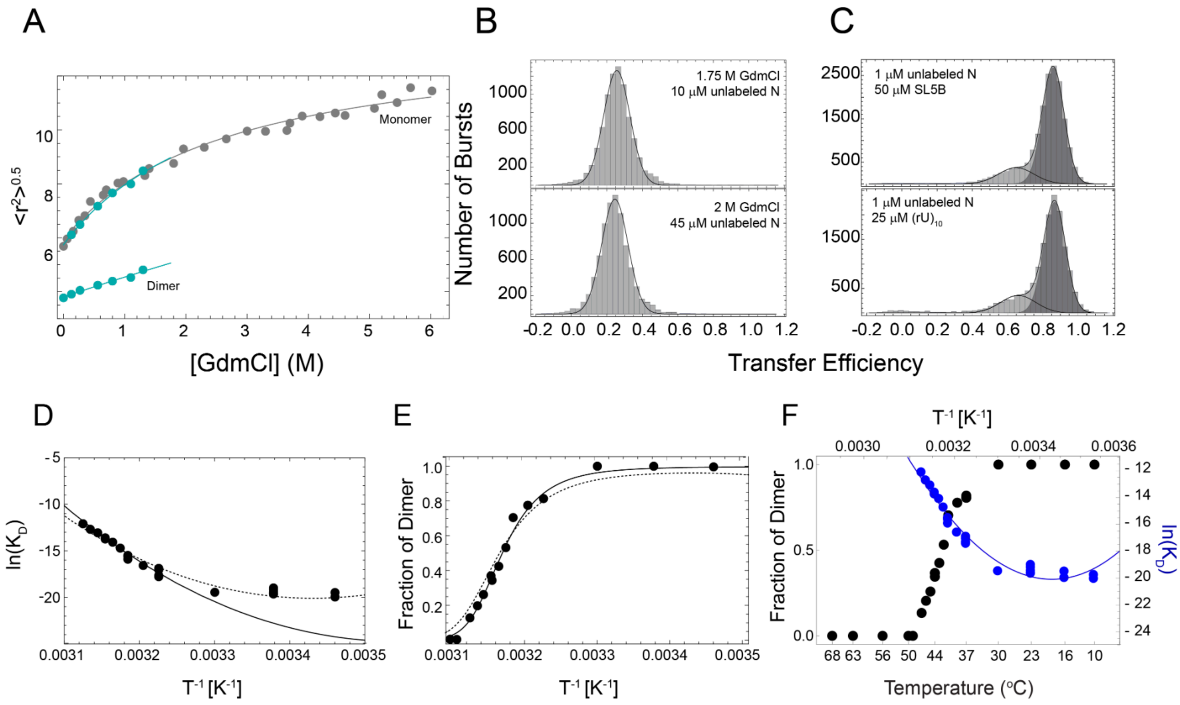


**Supplementary Figure 4. Effects on solution conditions on dimerization. A.** Root-mean-squared interdye distance for the $DD_{FL}$ as a function of denaturant. Grey points and fit represent the distance distributions for the $DD_{FL}$ at 100 pM total protein concentration (**Fig 2**). Teal points and fit represent the distance distributions for the $DD_{FL}$ at 1 μM total protein concentration (100 pM labeled + 1 μM unlabeled) (**Fig. 4**). **B.** Histogram of transfer efficiencies for 1.75 M GdmCl with the addition of 10 μM unlabeled (top) and 2 M GdmCl with the addition of 45 μM unlabeled (bottom). Extrapolation from the Van't Hoff dependence of $\ln(K_D)$ predicts a fraction of dimer of ~0.5; however, no dimer conformation is observed in either scenario. **C.** Histogram of transfer efficiencies at 37 °C (100 pM labeled + 1 μM unlabeled) with the addition of high concentrations of specific double-stranded RNA (top, SL5B) and non-specific single-stranded RNA (bottom, $(rU)_{10}$). **D-F.** Fit of the $\ln(K_D)$ as a function of the reciprocal of temperature (**D**) according to **Eq. 2b** (dashed line) compared to the fit of **Eq. 2b** to the fraction of the dimer population (**E**) (solid line). Overlap of the experimental trends and corresponding fit in panel **F**.

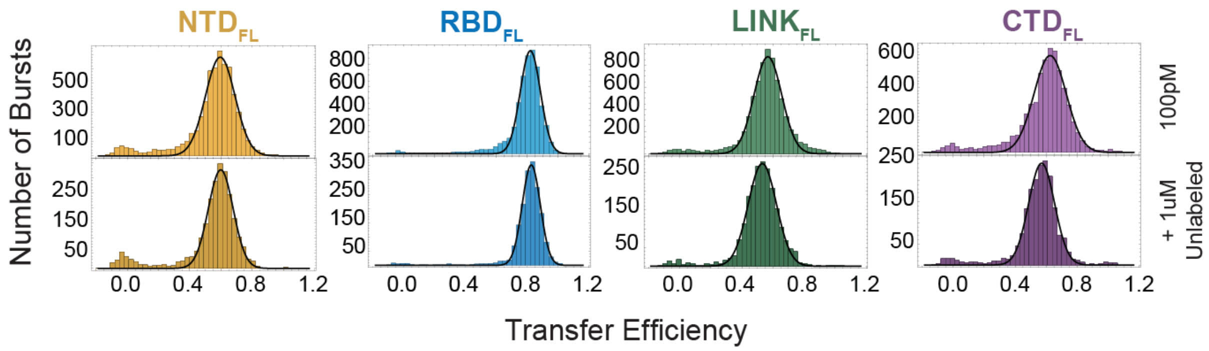


**Supplementary Figure 5. Dimerization alters the conformations of the Linker domain and CTD, but not NTD or RBD.** Transfer efficiency distribution for each labeled segment in the context of the full-length protein.

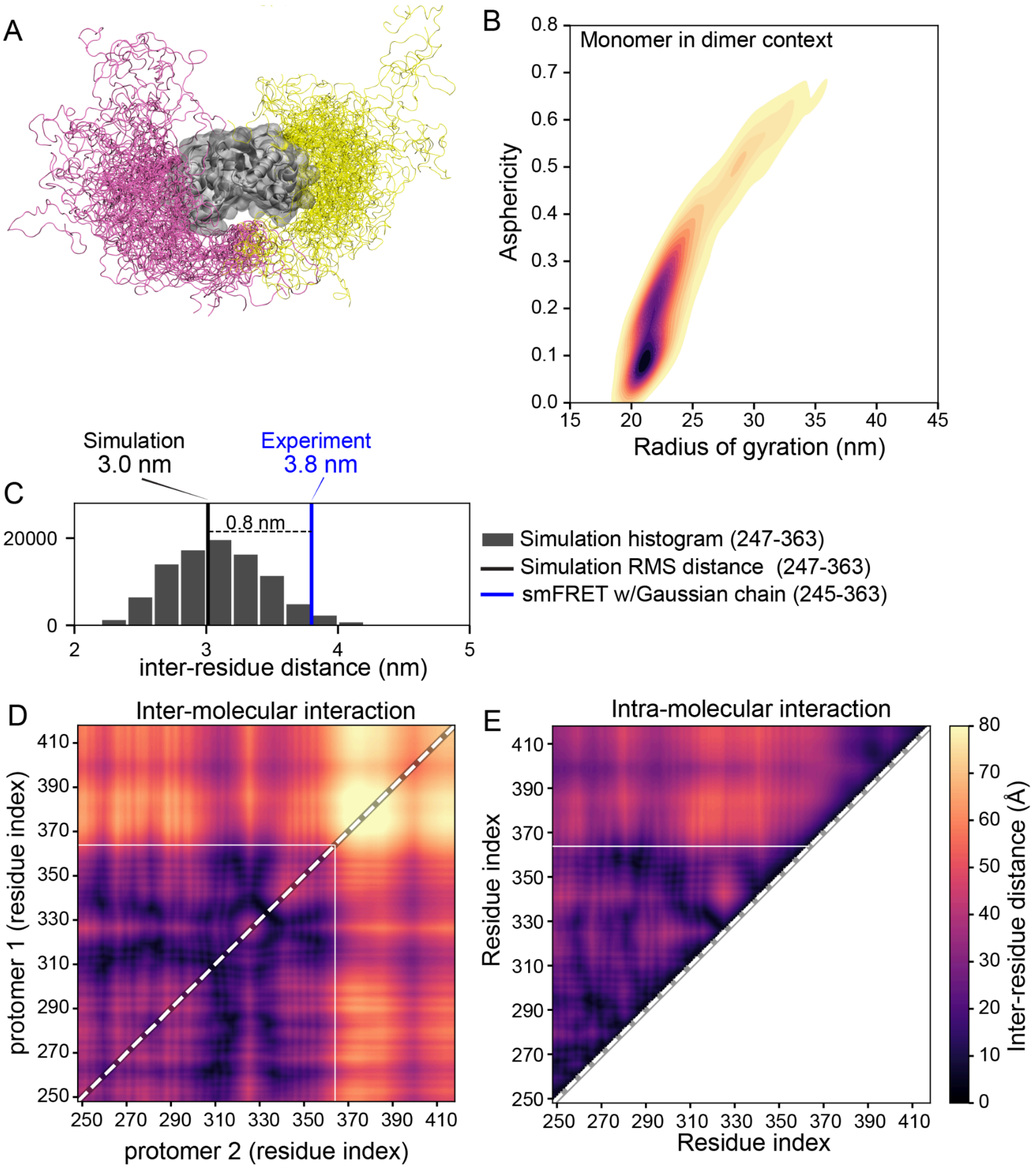


**Supplementary Figure 6. Simulations of the dimeric construct. A.** Rendering showing C-terminal IDRs superimposed over the dimeric structure. The IDRs largely do not interact with one another, but do transiently interact with the surface of the dimerization domain. **B.** Kernel density estimate showing the radius of gyration and asphericity for the monomer in the context of the dimer. Note the substantial reduction in average radius of gyration, asphericity, and the area adopted across the two-dimensional space (implicating a reduction in ensemble heterogeneity). **C.**

Comparison of root-mean-squared distance obtained from simulations and experiment. **D.** Inter-molecular distance map reporting on the average inter-molecular distance over the full ensemble. **E.** Intra-molecular distance map reporting on the average intra-molecular distance over the full ensemble (taken from both protomers).

#### Supplementary Tables

**Supplementary Table 1.** Sequence of Wildtype (WT) Nucleocapsid Protein. Labeling positions are reported as bold and underlined residues. Highlighted regions delineate folded domains.

| | |
|---|---|
| 1 | **M**SDNGPQNQR NAPRITFGGP SDSTGSNQNG ERSGARSKQR RPQGLPNNTA |
| 51 | SWFTALTQHG KEDLKFP**R**GQ GVPINTNSSP DDQIGYYRRA TRRIRGGDGK |
| 101 | MKDLSPRWYF YYLGTGPEAG LPYGANKDGI IWVATEGALN TPKDHIGTRN |
| 151 | PANNAAIVLQ LPQGTTLPKG F**Y**AEGSRGGS QASSRSSSRS RNSSRNSTPG |
| 201 | SSRGTSPARM AGNGGDAALA LLLLDRLNQL ESKMSGKGQQ QQGQ**T**VTKKS |
| 251 | AAEASKKPRQ KRTATKAYNV TQAFGRRGPE QTQGNFGDQE LIRQGTDYKH |
| 301 | WPQIAQFAPS ASAFFGMSRI GMEVTPSGTW LTYTGAIKLD DKDPNFKDQV |
| 351 | ILLNKHIDAY KT**F**PPTEPKK DKKKKADETQ ALPQRQKKQQ TVTLLPAADL |
| 401 | DDFSKQLQQS MSSADSTQ**A** |

**Supplementary Table 2. N protein constructs used in this study**

| Name | Sequence | Start Position (WT) | End Position (WT) | Labeling Positions |
|---|---|---|---|---|
| $NTD_{FL}$ | GPCSDNGPQNQRNAPRITFGGPSDSTGSN QNGERSGARSKQRRPQGLPNNTASWFTAL TQHGKEDLKFPCGQGVPINTNSSPDDQIG YYRRATRRIRGGDGKMKDLSPRWYFYYLG TGPEAGLPYGANKDGIIWVATEGALNTPK DHIGTRNPANNAAIVLQLPQGTTLPKGFY AEGSRGGSQASSRSSSRSRNSSRNSTPGS SRGTSPARMAGNGGDAALALLLLDRLNQL ESKMSGKGQQQQGQTVTKKSAAEASKKPR QKRTATKAYNVTQAFGRRGPEQTQGNFGD QELIRQGTDYKHWPQIAQFAPSASAFFGM SRIGMEVTPSGTWLTYTGAIKLDDKDPNF KDQVILLNKHIDAYKTFPPTEPKKDKKKK ADETQALPQRQKKQQTVTLLPAADLDDFS KQLQQSMSSADSTQA | 1 | 419 | M1C-R68C |
| $RBD_{FL}$ | GPMSDNGPQNQRNAPRITFGGPSDSTGSN QNGERSGARSKQRRPQGLPNNTASWFTAL TQHGKEDLKFPCGQGVPINTNSSPDDQIG | 1 | 419 | R68C-Y172C |

| | | | | |
|---|---|---|---|---|
| | YYRRATRRIRGGDGKMKDLSPRWYFYYLG<br>TGPEAGLPYGANKDGIIWVATEGALNTPK<br>DHIGTRNPANNAAIVLQLPQGTTLPKGFC<br>AEGSRGGSQASSRSSSRSRNSSRNSTPGS<br>SRGTSPARMAGNGGDAALALLLLDRLNQL<br>ESKMSGKGQQQQGQTVTKKSAAEASKKPR<br>QKRTATKAYNVTQAFGRRGPEQTQGNFGD<br>QELIRQGTDYKHWPQIAQFAPSASAFFGM<br>SRIGMEVTPSGTWLTYTGAIKLDDKDPNF<br>KDQVILLNKHIDAYKTFPPTEPKKDKKKK<br>ADETQALPQRQKKQQTVTLLPAADLDDFS<br>KQLQQSMSSADSTQA | | | |
| $LINK_{FL}$ | GPMSDNGPQNQRNAPRITFGGPSDSTGSN<br>QNGERSGARSKQRRPQGLPNNTASWFTAL<br>TQHGKEDLKFPRGQGVPINTNSSPDDQIG<br>YYRRATRRIRGGDGKMKDLSPRWYFYYLG<br>TGPEAGLPYGANKDGIIWVATEGALNTPK<br>DHIGTRNPANNAAIVLQLPQGTTLPKGFC<br>AEGSRGGSQASSRSSSRSRNSSRNSTPGS<br>SRGTSPARMAGNGGDAALALLLLDRLNQL<br>ESKMSGKGQQQQGQCVTKKSAAEASKKPR<br>QKRTATKAYNVTQAFGRRGPEQTQGNFGD<br>QELIRQGTDYKHWPQIAQFAPSASAFFGM<br>SRIGMEVTPSGTWLTYTGAIKLDDKDPNF | 1 | 419 | Y172C-<br>T245C |

| | | | | |
|---|---|---|---|---|
| | KDQVILLNKHIDAYKTFPPTEPKKDKKKK<br>ADETQALPQRQKKQQTVTLLPAADLDDFS<br>KQLQQSMSSADSTQA | | | |
| $DD_{FL}$ | GPMSDNGPQNQRNAPRITFGGPSDSTGSN<br>QNGERSGARSKQRRPQGLPNNTASWFTAL<br>TQHGKEDLKFPRGQGVPINTNSSPDDQIG<br>YYRRATRRIRGGDGKMKDLSPRWYFYYLG<br>TGPEAGLPYGANKDGIIWVATEGALNTPK<br>DHIGTRNPANNAAIVLQLPQGTTLPKGFY<br>AEGSRGGSQASSRSSSRSRNSSRNSTPGS<br>SRGTSPARMAGNGGDAALALLLLDRLNQL<br>ESKMSGKGQQQQGQCVTKKSAAEASKKPR<br>QKRTATKAYNVTQAFGRRGPEQTQGNFGD<br>QELIRQGTDYKHWPQIAQFAPSASAFFGM<br>SRIGMEVTPSGTWLTYTGAIKLDDKDPNF<br>KDQVILLNKHIDAYKTCPPTEPKKDKKKK<br>ADETQALPQRQKKQQTVTLLPAADLDDFS<br>KQLQQSMSSADSTQA | 1 | 419 | T245C-<br>F363C |
| $CTD_{FL}$ | GPMSDNGPQNQRNAPRITFGGPSDSTGSN<br>QNGERSGARSKQRRPQGLPNNTASWFTAL<br>TQHGKEDLKFPRGQGVPINTNSSPDDQIG<br>YYRRATRRIRGGDGKMKDLSPRWYFYYLG<br>TGPEAGLPYGANKDGIIWVATEGALNTPK<br>DHIGTRNPANNAAIVLQLPQGTTLPKGFY | 1 | 419 | F363C-<br>A419C |

| | | | | |
|---|---|---|---|---|
| | AEGSRGGSQASSRSSSRSRNSSRNSTPGS<br>SRGTSPARMAGNGGDAALALLLLDRLNQL<br>ESKMSGKGQQQQGQTVTKKSAAEASKKPR<br>QKRTATKAYNVTQAFGRRGPEQTQGNFGD<br>QELIRQGTDYKHWPQIAQFAPSASAFFGM<br>SRIGMEVTPSGTWLTYTGAIKLDDKDPNF<br>KDQVILLNKHIDAYKTCPPTEPKKDKKKK<br>ADETQALPQRQKKQQTVTLLPAADLDDFS<br>KQLQQSMSSADSTQC | | | |
| Single-Labeled$_{FL}$ | GPMSDNGPQNQRNAPRITFGGPSDSTGSN<br>QNGERSGARSKQRRPQGLPNNTASWFTAL<br>TQHGKEDLKFPRGQGVPINTNSSPDDQIG<br>YYRRATRRIRGGDGKMKDLSPRWYFYYLG<br>TGPEAGLPYGANKDGIIWVATEGALNTPK<br>DHIGTRNPANNAAIVLQLPQGTTLPKGFY<br>AEGSRGGSQASSRSSSRSRNSSRNSTPGS<br>SRGTSPARMAGNGGDAALALLLLDRLNQL<br>ESKMSGKGQQQQGQTVTKKSAAEASKKPR<br>QKRTATKAYNVTQAFGRRGPEQTQGNFGD<br>QELIRQGTDYKHWPQIAQFAPSASAFFGM<br>SRIGMEVTPSGTWLTYTGAIKLDDKDPNF<br>KDQVILLNKHIDAYKTCPPTEPKKDKKKK<br>ADETQALPQRQKKQQTVTLLPAADLDDFS<br>KQLQQSMSSADSTQA | 1 | 419 | F363C |

**Supplementary Table 3. FRETTraj parameters for Alexa 488 and Alexa 594.**

| FRETraj parameters |
|---|

```
{"Position":
        {"Alexa488":
           {"attach_id": 6,
            "linker_length": 12.5,
            "linker_width": 4.5,
            "dye_radius1": 5,
            "dye_radius2": 6,
            "dye_radius3": 3,
            "cv_fraction": 0.0,
            "cv_thickness": 6,
            "use_LabelLib": False},
         "Alexa594":
           {"attach_id": 936,
            "linker_length": 17,
            "linker_width": 3,
            "dye_radius1": 8,
            "dye_radius2": 5.7,
            "dye_radius3": 3,
            "cv_fraction": 0.0,
            "cv_thickness": 6,
            "use_LabelLib": False},
        },
      "Distance": {"Alexa488-Alexa594":
        {"R0": 54,
           "n_dist": 1000000}}
      }
```

**Supplementary Table 4. RNAs used in this study**

| RNA | Genomic position | nucleotides | Sequence 5'-3' | Origin |
|---|---|---|---|---|
| **Poly(rU)** | **-** | 10 | UUUUUUUUUU | IDT |
| **SL5B** | 228 - 252 | 30 | GGGCAUACCUAGGU UUCGUCCGGGUGU GCC | *in vitro transcribed* |
| **dsRNA$_{150F}$** | - | 150 | GGGATCTTGTGAGCG GATAACAATTATACTC TAGAAATAATTTTGTTT AACTTTAAGAAGGAG ATATATCATGAGCAGC AGCCATCATCATCATC ATCACAGCAGCGGCC GGTGAAGCGCGGCA GATATATTTGATGGGG TAAAATGGA | *in vitro transcribed and annealed* |
| **dsRNA$_{150R}$** | - | 150 | GGGTCCATTTTACCC CATCAAATATATCTGC CGCGCTTCACCAGGC CGCTGCTGTGATGAT GATGATGATGGCTGC TGCTCATGATATATCT CCTTCTTAAAGTTAAA CAAAATTATTTCTAGA GTATAATTGTTATCCG CTCACAAGAT | *in vitro transcribed and annealed* |

**Supplementary Table 5. Thermodynamic parameters fit from temperature dependence with 41 ℃ as reference temperature for dimerization of labeled N protein with unlabeled protein.**

| | **ΔH (kcal/mol)** | **ΔS (kcal/mol K)** | **$\Delta C_p$ (kcal/mol/K)** | **ΔG at 41 $^{O}$C (kcal/mol)** |
|---|---|---|---|---|
| Van't Hoff | - 99 ± 6 | - 0.29 ± 0.02 | - | -9 ± 7 |

| | **ΔH (kcal/mol)** | **ΔS (kcal/mol K)** | **$\Delta C_p$ (kcal/mol/K)** | **ΔG at 41 $^{O}$C (kcal/mol)** |
|---|---|---|---|---|
| fraction dimer | - 103 ± 5 | - 0.30 ± 0.02 | -3 ± 2 | -10 ± 7 |
| $\ln(K_D)$ | - 80 ± 5 | - 0.22 ± 0.02 | -3.5 ± 0.4 | -10 ± 6 |
